# Adaptive divergence in brain composition between ecologically distinct incipient species

**DOI:** 10.1101/043182

**Authors:** Stephen H. Montgomery, Richard M. Merrill

**Affiliations:** Dept. Genetics, Evolution & Environment, University College London, Gower Street, London, UK, WC1E 6BT; Dept. Zoology, University of Cambridge, Downing Street, Cambridge, UK, CB2 3EJ

## Abstract

During ecological speciation diverging populations are exposed to contrasting sensory and spatial information that present new behavioral and perceptive challenges. Here, we investigate how brain composition evolves during the early stages of speciation. The incipient species pair, *Heliconius erato cyrbia* and *H. himera*, have parapatric ranges across an environmental and altitudinal gradient. Despite continuing gene-flow, these species have divergent ecological, behavioral and physiological traits. We demonstrate that these incipient species also differ significantly in brain composition, especially in the size of sensory structures. *H. erato* has larger visual components whilst *H. himera* invests more heavily in olfaction. These differences are not explained by environmentally-induced plasticity, but reflect non-allometric shifts in brain structure. Our results suggest the adaptive evolution of brain structure and function play an important role in facilitating the emergence of ecologically distinct species, and imply that plasticity alone may be insufficient to meet the demands of novel environments.

## Introduction

Local adaptation following the colonization of novel environments promotes the origin of new species^1,2^. During the early stages of this process, diverging populations are exposed to contrasting sensory and spatial information that present new behavioral and perceptive challenges. These can be met by changes in brain function, often reflected in differential investment in brain components^3^. Analyses across phylogenetically disperse species suggest that adaptive changes in brain composition are driven by selection to meet the demands of a species’ ecology^4,5^. In contrast, recent intraspecific studies highlight the potential for neural plasticity to facilitate optimization of brain composition to local conditions ^6–9^ Little is known about the role of brain evolution and plasticity at the intersection of these evolutionary scales when new species emerge from locally specialized populations.

The role of plasticity during ecological speciation continues to be controversial^10^. Plasticity can increase fitness in new environmental conditions^7,8,11,12^, particularly after rapid environmental change^13^. Plasticity in trade-odds between sensory modalities could also facilitate rapid adaptation to contrasting niches without changing the energetic cost of sensory processing^14^. By promoting survival, adaptive plasticity could either facilitate speciation by enabling persistent exposure to contrasting environmental conditions, or inhibit speciation by facilitating local adaptation without the evolution of reproductive isolation^8,12,15^. Plasticity can also be costly^7^, particularly for energetically expensive neural tissue^16^. The benefits of plasticity may therefore be absent or transient depending on population dynamics and fitness landscape^17,18^. Indeed, plasticity may be maladaptive if populations are pushed further from fitness optima, increasing the strength of selection for heritable adaptations^15^. A shortage of case-studies has so far prohibited the resolution of this debate.

Here, we provide a novel case study focused on the roles of adaptation and plasticity in brain composition during the early stages of speciation in *Heliconius* butterflies. Speciation in *Heliconius* often involves selection favouring ecological divergence^19^ and a number of extant taxon-pairs provide ‘snap-shots’ of this process at different stages of completion^20,21^. We studied one example involving two incipient species of *Heliconius* butterfly that have recently diverged across an environmental gradient, and reflect the transition from polymorphic races to species; *H. himera* and *H. erato cyrbia. H. himera* is an incipient species emerging from within the *H. erato* clade^22^. Unlike low altitude races of *H. erato*, which are typically found in large-leaved secondary wet forest, *H. himera* is endemic to high altitude dry forest in the western border of Ecuador and Peru^23^. This parapatric distribution across an altitudinal gradient is maintained by strong selection^24^, and exposes individuals to different environmental conditions, including the distribution and intensity of different wavelengths of light, average rainfall and daily temperature range^25–27^. These contrasting abiotic conditions in turn shape differences in forest and foliage type, the ecological communities and predators individuals experience.

Adaptation to these contrasting environments has played a central role in driving speciation in these butterflies^24–27^. In *H. erato* and *H. himera*, strong divergent ecological selection, imposed by frequency-dependent predation of rare *Heliconius* warning patterns^28,29^, is augmented by assortative mating ^27,30,31^. Both migrant and hybrid individuals are thought to suffer fitness costs when poorly matched to their environment, due to behavioural or physiological divergence^27^. This ecological specialization in habitat preference persists despite ongoing gene flow across a narrow contact zone, and in which 5-10% of individuals are of hybrid origin^24,25^. High rates of hybridization emphasize the recent origin of *H. himera*. For example, the frequency of hybrids observed between the sympatric species pair *H. melpomene* and *H. cydno*, which diverged ~1 million years ago^32^, is less than 0.1%^33^.

Recently, the size of brain components in *Heliconius* have been shown to have significant amounts of ontogenetic and environmentally-induced plasticity^34^, which could potentially facilitate specialisation to contrasting habitats during the early stages of ecological speciation. Contrary to this hypothesis, we instead demonstrate that ecological divergence has favoured adaptive shifts in the relative size of multiple brain components between *H. himera* and *H. erato*. These species differences cannot be explained solely by environmentally induced plasticity, suggesting heritable adaptations in brain structure and function have contributed to the emergence of *H. himera*.

## Results and Discussion

We collected *H. himera* and *H. erato cyrbia* from the forests around Vilcabamba and Balsas Canton, respectively, in Southern Ecuador (Fig. 1). These populations lie either side of a narrow hybrid zone and have previously been studied to demonstrate divergence in habitat, ecology, behaviour and life history^25–27,31^. We quantified variation in the size of 13 major brain components, or ‘neuropils’, along with the remaining volume of the central brain (henceforth rCBR) from 16 individuals of each species using immunofluorescence staining and 3D image segmentation^35^. These include the major visual and olfactory neuropils, as well as the mushroom bodies, structures linked to learning and memory^36^, and components of the central complex, a multimodal integration and action selection center^37–39^.

**Figure 1:**
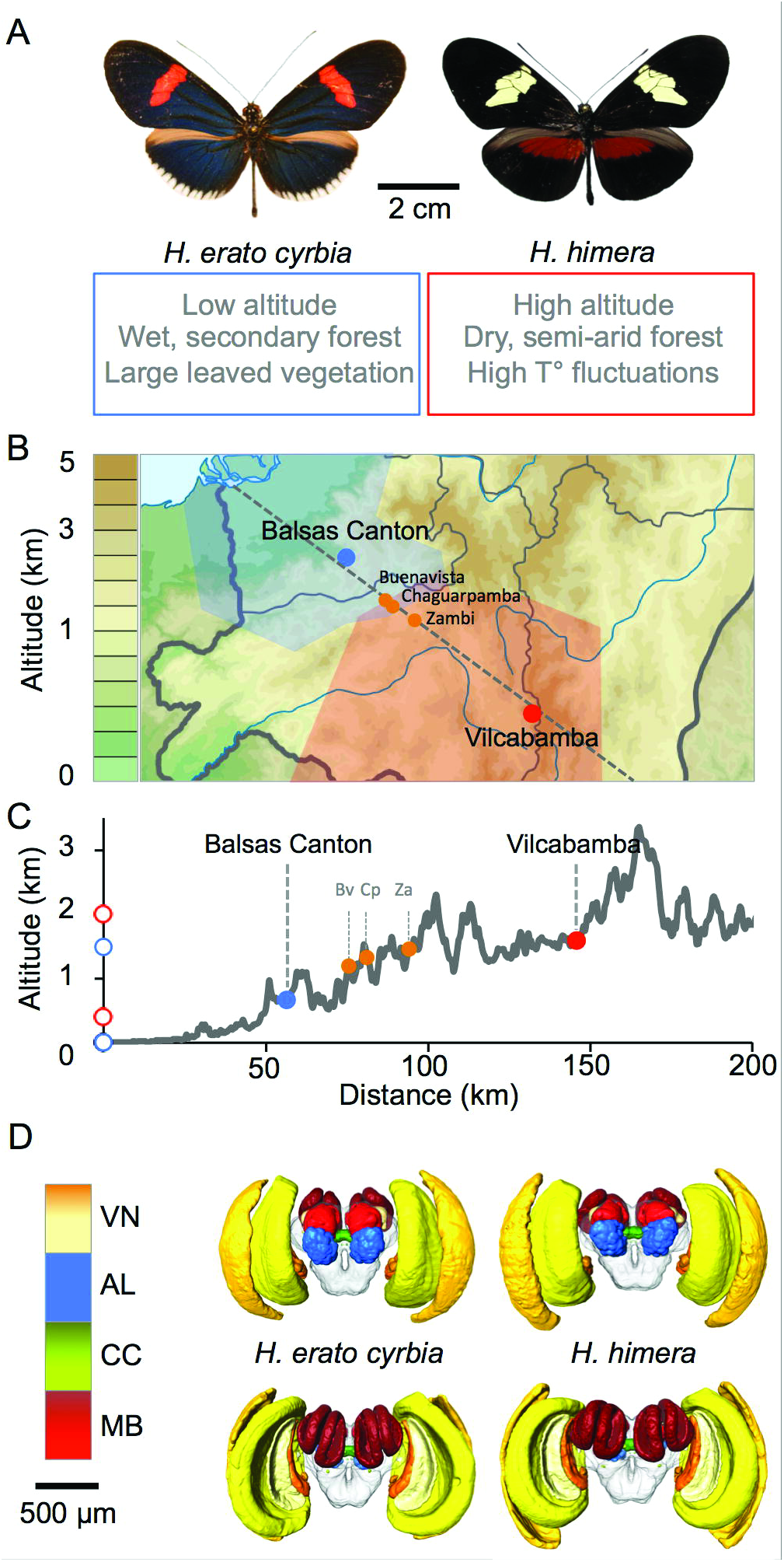
Distribution and ecology of *H. erato cyrbia* and *H. himera*. A) Colour patterns and key distinguishing features of the habitats of *H. e. cyrbia* (left) and *H. himera* (right). B) Approximate distribution of *H. e. cyrbia* (blue) and *H. himera* (red) in southern Ecuador showing the two sample localities, Balsas Canton and Vilcabamba, and the location of the hybrid zone south of Buenavista (Bv) and Chaguarpamba (Cp), but north of Zambi (Za). Geographic ranges are based on published data from Rosser et al.^57^ C) Variation in altitude across a transect illustrated by the grey dashed line in B. D) 3D volumetric models of the neuropil measured in *H. e. cyrbia* (left) and *H. himera* (right) viewed from the anterior (top row) and posterior (bottom row). VN: visual neuropil, AL: Antennal Lobe, CC: Central Complex, MB: Mushroom Body.

### Divergence in brain composition

Despite finding no significant differences between *H. himera* and *H. e. cyrbia* in the overall volume of the central brain (t_30_ = 0.688, p = 0.497) or total neuropil volume (t30 = 0.705, p = 0.487), a Principal Component Analysis (PCA) revealed marked divergence in brain composition between the two species. Using volumetric data for 13 neuropil and rCBR from 32 wild individuals, our PCA resulted in four major Principal Components (wPC), together explaining a total of 77.4% of the total variance. Of these, wPC2 (18.183% Var; F_1_ = 33.840, p < 0.001) and wPC4 (8.182% Var; F_1_ = 9.691, p = 0.004) were significantly associated with species identity (ANOVA controlling for sex) (Fig. 2A). This result is supported by a Discriminant Function Analysis (DFA), where the two species were separated along a single significant Discriminant Function (DF) (Wilks λ = 0.165, χ^2^ = 39.664, p < 0.001; Figure 2B) with 90% of individuals assigned to the correct species group with a high degree of confidence (p < 0.05). Re-analysis with four groups (species + sex), or on males and females separately, produced similar results (Supplemental Information). Across these multivariate analyses, components of the visual pathway including the medulla, lobula and lobula plate which are involved in processing of light, colour and movement, and the antennal lobe, the primary olfactory structure in the insect brain, had consistently strong contributions to PCs or DFs that separate *H. himera* and *H. e. cyrbia* (Supplemental Information).

**Figure 2:**
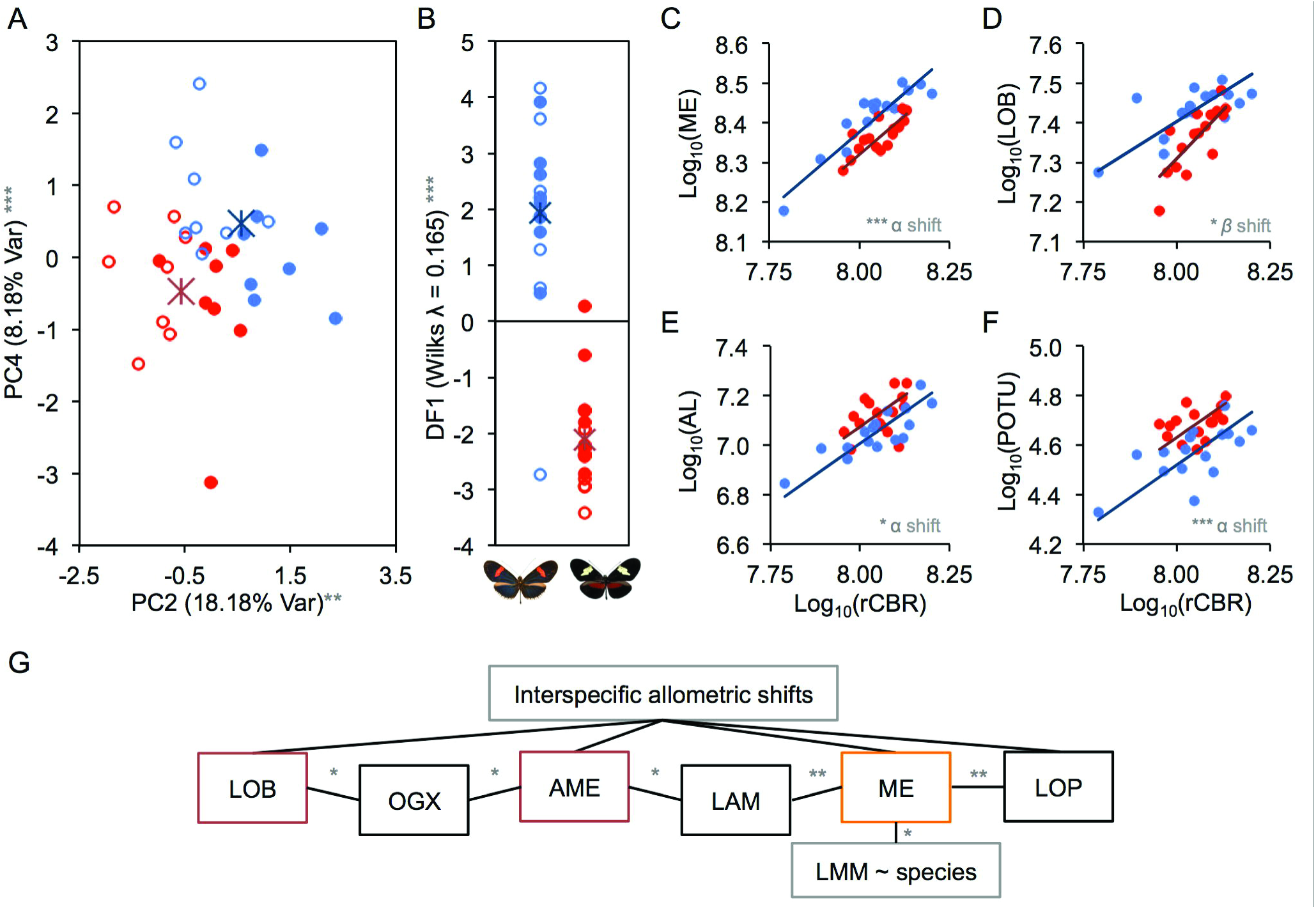
Divergence in brain composition in wild *H. e. cyrbia* and *H. himera*. A) Biplot between PC2 and PC4, which are both significantly different between species. *H. erato* are in blue, *H. himera* in red. Males are filled circles, females unfilled. B) Separation of species by Discriminant Function Analysis. Asterisks denote group means. C-F) Scaling relationships for all individuals between the central brain (rCBR) and the medulla (ME) (C), lobula (LOB) (D), antennal lobes (AL) (E) and posterior optic tubercule (POTU) (F). G) Patterns of covariance between neuropils in the optic lobe. Significant covariance is shown by solid black lines, with those neuropil with significantly different scaling relationships with rCBR between species shown above (interspecific allometric shifts), and those associated with species in the multiple regression shown below (LMM ~ species). * p < 0.05, ** p < 0.01, *** p < 0.001.

To further explore how individual neuropil contribute to species differences in brain composition, we next examined the scaling relationship between each neuropil and an independent measure of overall brain size (the unsegmented central brain; rCBR). The standard allometric scaling relationship (log *y* = *β* log *x* + *α*) provides a means to test for significant shifts in the allometric exponent (*β*) or scaling factor (*α*) between species, which together describe the relationship between two traits. Conserved scaling relationships are typically interpreted as indicating the presence of some constraint that results in covariance between variables. This constraint may arise from shared developmental mechanisms (or pleiotropy), or be due to selective covariance to maintain a constant level of functional integration^40^. Deviation from a shared scaling relationship can therefore indicate an adaptive change in the functional relationship between two phenotypes^41^.

The majority of neuropil in the optic lobes display non-allometric shifts in scaling with rCBR between species (Supplementary Information). After correcting for multiple tests using the false-discovery rate^42^ (for 13 neuropils), both the medulla (FDR-p < 0.001) and lobula plate (FDR-p = 0.026) show significant grade-shifts between species (difference in *α*), whilst the lobula shows a species difference in *β*(FDR-p = 0.026) (Fig. 2C-D). There is also some support for the accessory medulla displaying a grade-shift (nominal-p = 0.044). In all cases these differences result in an increase in the size of these structures in *H. e. cyrbia*. In contrast, we identified two central brain neuropil, the antennal lobe (FDR-p = 0.042) and the posterior optic tubercule (FDR-p < 0.001), which show grade-shifts towards larger volumes in *H. himera* (Fig. 2E-F). These differences in scaling relationships reflect substantial differences in volume. For example, for a given brain volume the medulla and lobula will be 12.3% and 18.2% larger in *H. e. cyrbia* respectively, whilst in *H. himera* the antennal lobe will be 14.5% larger and the posterior optic tubercule 22.6% larger.

The results of our scaling analyses are largely consistent regardless of whether sexes are pooled or considered separately, or whether rCBR or an alternative measure of overall size (total neuropil minus the neuropil of interest) is used (Supplemental Information). Importantly, because neither rCBR nor total brain size vary between species, these differences represent changes in the volume of individual neuropil, not a concerted size change affecting all neuropil, or a shift in rCBR volume.

### Covariance and composite adaptations

Our analyses demonstrate that at least three of the six neuropil in the optic lobes are larger in *H. erato*. These neuropil process visual information and are both functionally interdependent and physically connected by projection neurons^43,44^. It is therefore likely that if one component expands, this would have knock-on effects on other neuropils. We analysed patterns of covariance between visual neuropils using linear multiple regressions (controlling for species and sex) to assess whether the change in scaling relationships for multiple individual optic lobe neuropils reflect independent adaptations. This revealed that the six neuropil form a co-varying network (Fig. 2G), partially reflecting patterns of connectivity^43^. Correcting for this covariance, the association with species only remains significant for the medulla. This suggests changes in medulla size may be driving changes elsewhere in optic lobe. For example, medulla volume strongly co-varies with lobula plate volume (p = 0.003), but also shows some evidence of co-variance with lobula volume (p = 0.069; Supplemental Information).

We further investigated this possibility by examining the pairwise scaling relationships between medulla, lobula, lobula plate and accessory medulla. Consistent with the conclusion that variation in the size of the medulla drives changes in lobula plate size these neuropil show a major-axis shift between species along a conserved scaling relationship (Wald χ^2^ = 5.105, p = 0.024). However, there is also evidence of species differences in scaling exponent between the lobula and both the medulla and lobula plate (Likeihood Ratio = 12.275, p < 0.001 and LR = 5.039, p = 0.025 respectively). The accessory medulla volume shows a grade-shift in scaling with the medulla, lobula and lobula plate, consistent with a lesser effect of species identity on this neuropil (all p < 0.001; Supplemental Information). These analyses suggest that the species differences in size of the medulla and lobula plate may constitute a concerted adaptation, maintaining but expanding their functional relationship, whilst altering their functional association with the lobula.

We identify one co-varying network amongst components of the central brain; between antennal lobe volume, the mushroom body lobes and the mushroom body calyx. This may reflect the well-established role of the mushroom bodies in olfactory learning^45^. We found no significant association between antennal lobe and posterior optic tubercule volume, or between either of these neuropils and medulla (Supplementary Information), indicating that these may reflect functionally independent adaptations.

### Plasticity does not explain species differences

Plasticity in the neural development and behavior has been implicated in facilitating local adaptation and catalyzing speciation by promoting survival in novel habitats^7,8,18,46^. Several recent studies suggest plasticity in the development of brain composition contributes to locally adapted ecological morphs within species^6,9^. However, plasticity has significant costs^7^ and the net benefits may therefore be transitory^18^.

*Heliconius* brains show significant amounts of environment-dependent and independent post-eclosion growth^34^. To test whether plasticity explains the differences we observe between *H. erato* and *H. himera*, we obtained data for an additional 10 individuals of each species reared in a common environment and repeated the analyses described above. These individuals were the progeny of females collected at the same populations as our wild individuals. First, a PCA of all neuropil volumes separated the variance across 4 PCs (iPC). Of these, iPC1 (35.068% Var, F_1_ = 9.887, p = 0.006) and iPC2 (17.672% Var, F_1_ = 17.672, p = 0.001) were significantly associated with species identity. Similar results were obtained when wild and insectary reared individuals were analysed together (Fig. 3; Supplemental Information).

**Figure 3:**
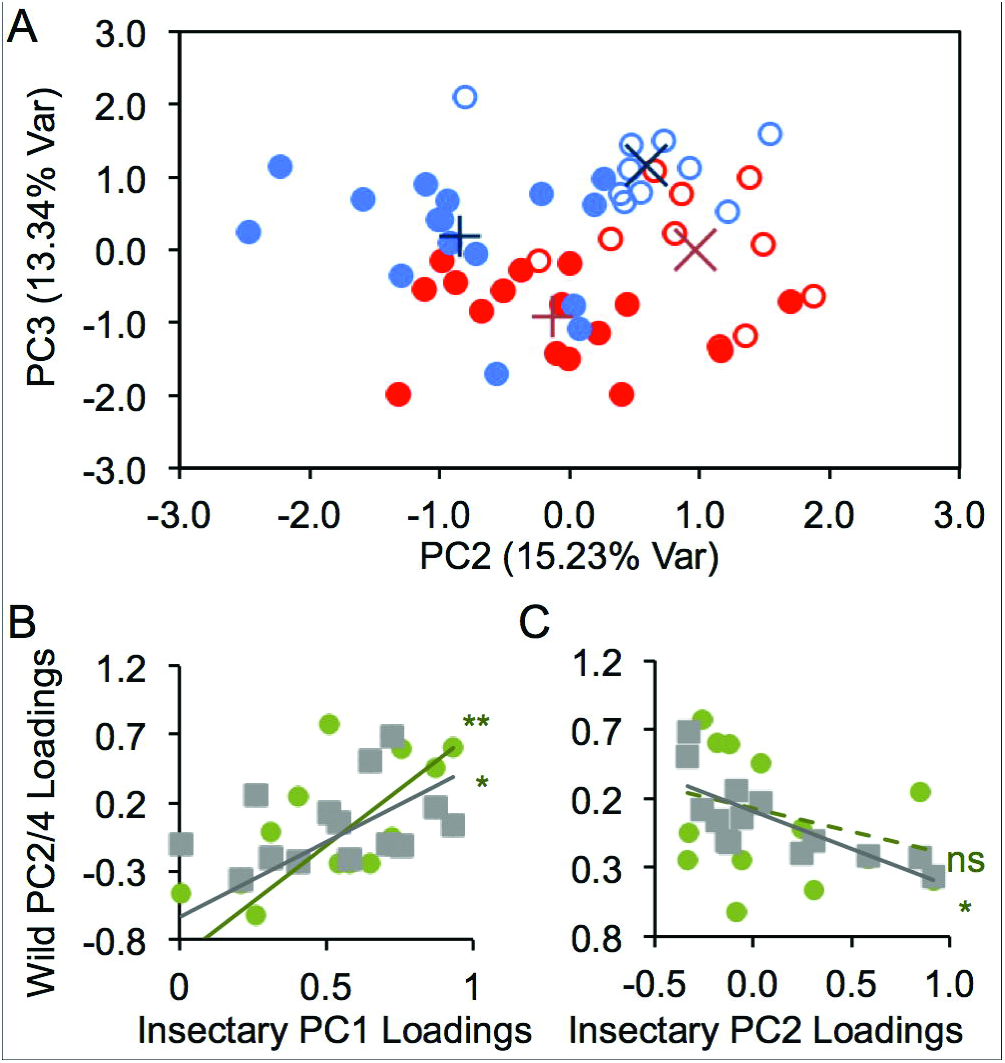
Consistent signal of divergence between wild and reared individuals. A) Biplot of PC2 and PC3 from a pooled analysis of wild (filled circles) and reared individuals (unfilled circles). *H. erato* are in blue, *H. himera* in red. Both PC2 (15.230%; Var; F_1_ = 70.670, p <0.001) and PC3 (13.336%; Var; F_1_ = 26.384, p <0.001) show significant differences between species. Asterisks denote group means. B-C) Regressions between the loadings of the measured neuropil on iPCs explaining species differences in reared individuals (x-axis) and wPCs explaining species differences in wild individuals (y-axis; wPC2 = green, wPC4 = grey).

We assessed whether the neuropils contributing to these iPCs were the same as those contributing to wPC2 and wPC4 in the wild caught samples by using a regression analysis of the loading coefficients for each neuropil. Loadings of neuropil on iPC1 from the insectary-reared analysis were significantly associated with loadings on both PCs associated with species from the wild analyses (wPC2: t_9_ = 3.438, p = 0.007; wPC4: t_9_ = 2.440, p = 0.037). Loadings on iPC2 are also significantly associated with loadings on wPC4 (wPC2: t_9_ = −1.309, p = 0.223; wPC4: t_9_ = −3.223, p = 0.001). Neither iPC1 or iPC2 show any association with wPC1 or wPC3 which do not vary between species (all p > 0.100). A DFA also shows strong support for species differences (Wilks λ = 0.028, χ^2^ = 39.456, p < 0.001) and correctly assigns 100% of insectary-reared individuals to the correct species group with a high degree of confidence (p < 0.001). The DF coefficients again implicate the visual neuropil and posterior optic tubercule as potential contributors to this difference (Supplemental Information). Together these collective results strong imply that the relative contribution of each neuropil to the species differences in brain composition in the comparison between insectary-reared individuals is similar to that between wild individuals.

Further analyses of the scaling relationships between each neuropil and rCBR largely confirm this conclusion. We identify the same grade-shifts towards larger volumes in *H. e. cyrbia* in medulla, lobula and lobula plate, and also in two further neuropil in the optic lobes; the lamina and ventral lobula (all FDR-p < 0.05; Supplemental Information). However, grade-shifts against central brain volume are not found for the antennal lobe or posterior optic tubercule. Although these are recovered using total neuropil volume as an alternative variable, this may indicate some contribution of species differences in plasticity to the results uncovered in wild individuals (Supplemental Information).

### Divergence in neuropil volume suggests a shift in the importance of sensory modalities

Our results imply a shift in investment in different sensory modalities in *H. e. cyrbia* and *H. himera* that may relate to their preferred habitat type. *H. e. cyrbia* invests in larger visual neuropil than *H. himera*, most notably the medulla, lobula and lobula plate. These neuropil have specific roles in processing visual information. In other insects the medulla plays a role in the parallelization of photoreceptor signals^43^ but also contains many processing elements with dual roles in colour-vision and motion detection pathways^47–49^. The lobula and lobula plate integrate visual information to extract abstract features such as shape and motion; for example the lobula plate is the primary site for motion computation in insects and tracking in the optic lobe, whilst the lobula has been linked to escape and chasing behavior^50^. Notably, the cellular architecture of the lobula plate is known to vary extensively across species in association with differences in flight behavior^51^. Given the difference in forest type inhabited by *H. himera* and *H. e. cyrbia* the volumetric difference in these components may reflect contrasting demands of visual and/or spatial information related to the density of vegetation or leaf size, and subsequent changes in light intensity and polarization.

In contrast, *H. himera* has higher levels of relative investment in the primary olfactory neuropil, the antennal lobe. Antennal lobes are comprised of glomeruli that are innervated by axons from olfactory sensory neurons in the antennae, which expresses olfactory receptors^52^. These glomeruli are arranged around the antennal lobe hub where information from different glomeruli are integrated to create a combinatorial code for odour cues^53^. The number of glomeruli is relatively constant across Lepidoptera^54^, including *Heliconius*^34^ Across more distantly related butterflies variation in antennal lobe size is disproportionately due to variation in the volume of the antennal lobe hub^34^. This suggests the complexity of neuronal branching in the antennal lobe hub, which likely reflects the complexity of the combinatorial code, dominates over changes in odour sensitivity, which is reflected by the volume of glomeruli^34^. Relative to central brain size, both the antennal lobe glomeruli (Wald χ^2^ = 5.674, p = 0.017) and hub (Wald χ^2^ = 11.106, p < 0.001) are expanded in wild *H. himera*, whilst maintaining a constant scaling relationship (*β-shift* LR < 0.001, p = 0.991; *α-shift* Wald χ^2^ = 0.940, p = 0.330). This may suggest the foraging or reproductive behavior of *H. himera* has a greater reliance on olfactory sensitivity, without changes in the complexity of olfactory repertoire utilized. The second striking expansion in *H. himera* is in one of the smallest components of the central complex, the posterior optic tubercule. In other insects, this neuropil receives a variety of inputs, including visual information from the accessory medulla, as well as mechanosensory and chemosensory information from the antennal lobes and other body parts^37^. Although we did not find statistical support for covariation in antennal lobe and posterior optic tubercule volume, the expansion of the posterior optic tubercule could conceivably reflect an increased input from the antennal lobe.

Finally, we note that our results mirror those found across more distantly related Lepidoptera with more extreme differences in ecology. For example, nocturnal moths and diurnal butterflies can be distinguished on the basis of differential expansion of the antennal lobe or medulla and lobula system^34,54^. Similarly, the Neotropical diurnal butterfly *Godyris zavaleta*, which is found in dark inner-forest has increased investment in the antennal lobe relative to *Heliconius* or *Danaus* which occupy habitats with greater light intensity^34,54^. This suggests similar selective pressures associated with divergent sensory environments may be shaping Lepidopteron brain composition across short and long evolutionary time-scales.

### Conclusion

Speciation across environmental gradients demands local adaptation to distinct environments^1,2^. By focusing of a pair of incipient species we have demonstrated that this exerts selective pressure on brain composition, resulting in significant non-allometric shifts in a specific suite of brain components. Under the assumption that scaling relationships reflect stabilising selection to maintain developmental or functional associations, these non-allometric changes are likely to be driven by selection for adaptive divergence, rather than being the result of phenotypic drift.

Although plasticity may facilitate ecological divergence initially^6–9^, especially where continued gene flow prevents the build up of adaptive alleles, the costs of plasticity are predicted to render this a transitory phase^7,18^. Our results demonstrate that even at the early stages of speciation, where gene flow persists^22,24^, plasticity alone cannot explain these species differences. We suggest selection on brain structure and function may commonly play a role in facilitating the early stages of ecological speciation, and that heritable divergence will quickly outweigh the contribution of plasticity.

## Experimental Procedures

### Animals

Brains of wild-caught individuals were fixed within 5 hours of collection, sampling eight individuals of each sex for both species. Insectary-reared individuals were bred from wild-caught parents and raised on a common hostplant (*Passiflora biflora*) in controlled conditions in cages (c. 1 × 2 × 2 m) of mixed sex and equal densities. Five individuals of each sex were sampled for both species, aged to 10-14 days, when both sexes are sexually mature.

### Immunocytochemistry and imaging

Brains were fixed in-situ using a Zinc-Formaldehyde solution, following a published protocol^35^. Further methodological details and anatomical descriptions of the *Heliconius* brain are available in Montgomery et al.^34^. Neuropil structure was revealed using immunofluorescence staining against a vesicle-associated protein at presynaptic sites, synapsin (anti-SYNORF1; obtained from the Developmental Studies Hybridoma Bank, University of Iowa, Department of Biological Sciences, Iowa City, IA 52242, USA; RRID: AB_2315424) and Cy2-conjugated affinity-purified polyclonal goat anti-mouse IgG (H+L) antibody (Jackson ImmunoResearch Laboratories, West Grove, PA), obtained from Stratech Scientific Ltd., Newmarket, Suffolk, UK (Jackson ImmunoResearch Cat No. 115-225-146, RRID: AB_2307343).

All imaging was performed on a confocal laser-scanning microscope (Leica TCS SP8, Leica Microsystem, Mannheim, Germany) using a 10× dry objective with a numerical aperture of 0.4 (Leica Material No. 11506511), a mechanical *z*-step of 2 μm and an *x-y* resolution of 512 × 512 pixels. The *z*-dimension was scaled 1.52× to correct the artifactual shortening^34^. We assigned image regions to brain components using the Amira 5.5 *labelfield* module and defining outlines based on the brightness of the synapsin immunofluorescence. We reconstructed total central brain volume (CBR),six paired neuropils in the optic lobes, six paired and one unpaired neuropils in the central brain (CBR), and measured their volume using the *measure statistics* module. The total volume of segmented structures in the CBR was subtracted from total CBR volume to obtain a measure of unsegmented CBR (rCBR). Due to the lack of volumetric asymmetry in *Heliconius* neuropil^34^ we measured the volume of paired neuropil from one hemisphere, chosen at random, and multiplied the measured volume by two.

### Statistical analyses

Multivariate analyses were performed in SPSS v. 22 (SPSS Inc., Chicago, IL). Principal Component Analyses (PCA) were performed using segmented structures and rCBR. Species differences in PC values were analyzed using an ANOVA, including species and sex as binary cofactors, in R^55^. We complemented this analysis with a Discrimant Function Analysis (DFA) to test how reliably individuals can be assigned to their respective groups on the basis of their volumetric differences in neuropil. In this analysis, Wilks’ λ provides a measure of the proportion of total variance not explained by group differences, and the χ^2^ statistic provides a test for significant group differences.

We further explored whether the scaling relationships between each component and rCBR were conserved across *H. himera* and *H. erato cyrbia* using major axis regressions in SMATR v.3.4-3^56^. Using the standard allometric scaling relationship: log *y* = *β* log *x* + *α*, we performed tests for significant shifts in the allometric slope (*β*) between the species. This was followed by two further tests which assume a common slope: 1) for differences in *α* that suggest discrete ‘gradeshifts’ in the relationship between two variables, 2) for major axis-shifts along a common slope. Covariance between neuropils was investigated using multiple regression. All volumes were log_10_-transformed before data analysis.

**Table 1.**
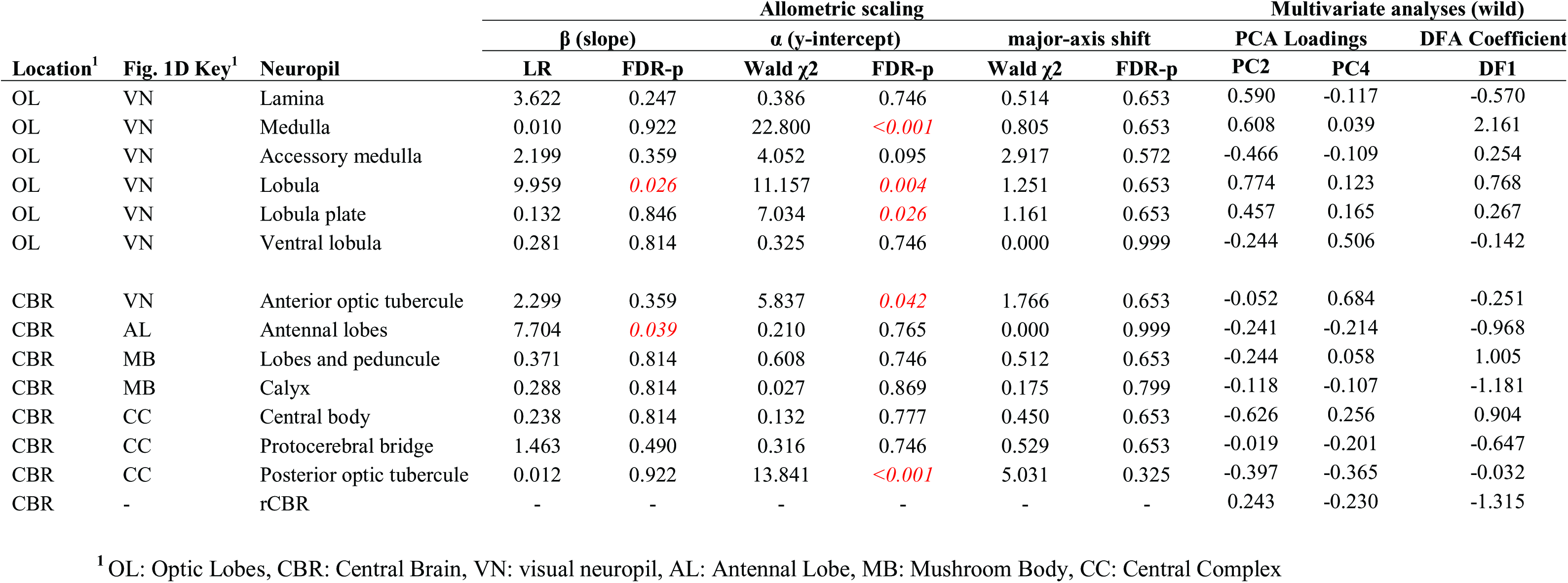

| Location<sup>1</sup> | Fig. 1D Key<sup>1</sup> | Neuropil | Allometric scaling |  |  |  |  |  | Multivariate analyses (wild) |  |  |
| --- | --- | --- | --- | --- | --- | --- | --- | --- | --- | --- | --- |
| | | | $\beta$ (slope) | | $\alpha$ (y-intercept) | | major-axis shift | | PCA Loadings | | DFA Coefficient |
| | | | LR | FDR-p | Wald $\chi^2$ | FDR-p | Wald $\chi^2$ | FDR-p | PC2 | PC4 | DF1 |
| OL | VN | Lamina | 3.622 | 0.247 | 0.386 | 0.746 | 0.514 | 0.653 | 0.590 | -0.117 | -0.570 |
| OL | VN | Medulla | 0.010 | 0.922 | 22.800 | <0.001 | 0.805 | 0.653 | 0.608 | 0.039 | 2.161 |
| OL | VN | Accessory medulla | 2.199 | 0.359 | 4.052 | 0.095 | 2.917 | 0.572 | -0.466 | -0.109 | 0.254 |
| OL | VN | Lobula | 9.959 | 0.026 | 11.157 | 0.004 | 1.251 | 0.653 | 0.774 | 0.123 | 0.768 |
| OL | VN | Lobula plate | 0.132 | 0.846 | 7.034 | 0.026 | 1.161 | 0.653 | 0.457 | 0.165 | 0.267 |
| OL | VN | Ventral lobula | 0.281 | 0.814 | 0.325 | 0.746 | 0.000 | 0.999 | -0.244 | 0.506 | -0.142 |
| CBR | VN | Anterior optic tubercle | 2.299 | 0.359 | 5.837 | 0.042 | 1.766 | 0.653 | -0.052 | 0.684 | -0.251 |
| CBR | AL | Antennal lobes | 7.704 | 0.039 | 0.210 | 0.765 | 0.000 | 0.999 | -0.241 | -0.214 | -0.968 |
| CBR | MB | Lobes and peduncle | 0.371 | 0.814 | 0.608 | 0.746 | 0.512 | 0.653 | -0.244 | 0.058 | 1.005 |
| CBR | MB | Calyx | 0.288 | 0.814 | 0.027 | 0.869 | 0.175 | 0.799 | -0.118 | -0.107 | -1.181 |
| CBR | CC | Central body | 0.238 | 0.814 | 0.132 | 0.777 | 0.450 | 0.653 | -0.626 | 0.256 | 0.904 |
| CBR | CC | Protocerebral bridge | 1.463 | 0.490 | 0.316 | 0.746 | 0.529 | 0.653 | -0.019 | -0.201 | -0.647 |
| CBR | CC | Posterior optic tubercle | 0.012 | 0.922 | 13.841 | <0.001 | 5.031 | 0.325 | -0.397 | -0.365 | -0.032 |
| CBR | - | rCBR | - | - | - | - | - | - | 0.243 | -0.230 | -1.315 |

<sup>1</sup> OL: Optic Lobes, CBR: Central Brain, VN: visual neuropil, AL: Antennal Lobe, MB: Mushroom Body, CC: Central Complex

## Author Contributions

Study conception and fieldwork: SHM and RMM. Insectary rearing: RMM. Dissections, acquisition of image data, analysis, interpretation and initial manuscript draft: SHM. Final interpretation and drafting: SHM and RMM.

## Acknowledgements

We thank the Ministerio del Ambiente del Ecuador for permission to collect butterflies. SHM was supported by research fellowships from the Royal Commission for the Exhibition of 1851 and the Leverhulme Trust, a Royal Society Research Grant (RG110466) and a British Ecological Society Early Career Project Grant. RMM was supported by a Junior Research Fellowship at King’s College, Cambridge, and funding from the Bedford Fund. We are also grateful to Chris Jiggins for providing insectary space and Swidbert Ott for discussions on insect brain evolution.

